# The *Wolbachia* CinB Nuclease is Sufficient for Induction of Cytoplasmic Incompatibility

**DOI:** 10.1101/2021.10.22.465375

**Authors:** Guangxin Sun, Mengwen Zhang, Hongli Chen, Mark Hochstrasser

## Abstract

*Wolbachia* are obligate intracellular bacteria that can alter reproduction of their arthropod hosts, often through a mechanism called cytoplasmic incompatibility (CI). In CI, uninfected females fertilized by infected males yield few offspring, but if both are similarly infected, normal embryo viability results (called ‘rescue’). CI factors (Cifs) responsible for CI are pairs of proteins encoded by linked genes. The downstream gene in each pair encodes either a deubiquitylase (Ci<u>d</u>B) or a nuclease (Ci<u>n</u>B). The upstream gene products, CidA and CinA, bind their cognate enzymes with high specificity. Expression of CidB or CinB in yeast inhibits growth, but growth is rescued by expression of the cognate CifA protein. By contrast, transgenic *Drosophila* male germline expression of both *cifA* and *cifB* was reported to be necessary to induce CI-like embryonic arrest; *cifA* expression alone in females is sufficient for rescue. This pattern, seen with genes from several *Wolbachia* strains, has been called the ‘2-by-1’ model. Here we show male germline expression of the *cinB* gene alone, from a distinct clade of *cif* genes from *w*No *Wolbachia*, is sufficient to induce nearly complete loss of embryo viability. This male sterility is fully rescued by cognate *cinA*^*w*No^ expression in the female germline. The proteins behave similarly in yeast. CinB^*w*No^ toxicity depends on its nuclease active site. These results demonstrate that highly divergent CinB nucleases can induce CI, that rescue by cognate CifA factors is a general feature of *Wolbachia* CI systems, and that CifA is not required in males for CI induction.

**Importance:** *Wolbachia* are bacteria that live within the cells of many insects. Like mitochondria, they are only inherited from females. *Wolbachia* often increase the number of infected females to promote spread of infection using a type of male sterility called cytoplasmic incompatibility (CI): when uninfected females mate with infected males, most embryos die; if both are similarly infected, embryos develop normally, giving infected females an advantage in producing offspring. CI is being used against disease-carrying mosquitoes and agricultural pests. *Wolbachia* proteins called CifA and CifB, which bind one other, cause CI, but how they work has been unclear. Here we show that a CifB protein singly produced in fruit fly males causes sterility in crosses to normal females, but this is rescued if the females produce the CifA partner. These findings clarify a broad range of observations on CI and will allow more rational approaches to using it for insect control.

## Introduction

Bacteria-arthropod symbioses are extremely common and range from full parasitism to mutualism (1). Possibly the most successful bacterial endosymbiont in the world is the obligate intracellular α-bacterium *Wolbachia pipientis*, which infects ~40% of all terrestrial arthropod species (2). *Wolbachia* is best known for its ability to manipulate the reproduction of its hosts in ways that increase inheritance of the bacteria through the female germline (3). Manipulations include parthenogenesis, male killing, and feminization of chromosomal males, but the most common is a mechanism called cytoplasmic incompatibility or CI (4). In CI, when females are infected, fertilization by either infected or uninfected males yields normal numbers of viable embryos, but if an uninfected female mates with a *Wolbachia-infected* male, a high fraction of the resulting embryos die. The ability of *Wolbachia*-infected females to ‘rescue’ viability provides a selective advantage to infected females and can drive *Wolbachia* infections into populations. Several other bacterial species are now known to cause CI, but *Wolbachia* is the most widespread and best studied (5).

*Wolbachia*-induced CI is being used in several ways to control mosquito-vectored disease, particularly by *Aedes aegypti* mosquitoes that transmit dengue virus and other arboviruses. In one approach, introduction of large numbers of *Wolbachia*-infected male mosquitoes into uninfected mosquito populations causes massive drops in mosquito number due to the post-zygotic male sterility caused by CI (6). Another control method exploits the observation that *Wolbachia* infection severely limits the ability of insects to carry certain arboviruses. In this approach, male and female *A. aegypti* mosquitoes trans-infected with *Wolbachia*, usually the *w*Mel strain, are released together in areas with endemic arbovirus-infected mosquitoes–principally those carrying dengue virus (7, 8). Due to the ability of the *w*Mel strain to cause CI, the bacterial infection rapidly spreads, and dramatic reductions in local dengue fever cases have been reported (7, 9).

The molecular basis of CI had long been a mystery, but recently, the *Wolbachia* factors responsible were identified (10–12). The CI factors (Cifs) are encoded by two-gene operons, with the upstream gene designated most generally as *cifA* and the downstream gene as *cifB* (see Figure 2A). Different CI-inducing *Wolbachia* strains carry distinct but related *cifA-cifB* operons, and some strains bear multiple divergent versions, which in some cases involve pseudogenes (13, 14). The downstream gene in each cognate gene pair usually encodes either a deubiquitylase (DUB) that cleaves ubiquitin from substrate proteins (CI-inducing DUB or CidB) or a DNA-cleaving enzyme (CI-inducing nuclease or CinB) (15, 16). The *w*Pip strain has syntenic gene pairs of both types. When either CidB^*w*Pip^ or CinB^*w*Pip^ is expressed by itself in the yeast *Saccharomyces cerevisiae*, temperature-dependent growth inhibition is observed (10). Growth impairment depends on the catalytic activity of the respective enzymes.

**Figure 1.**
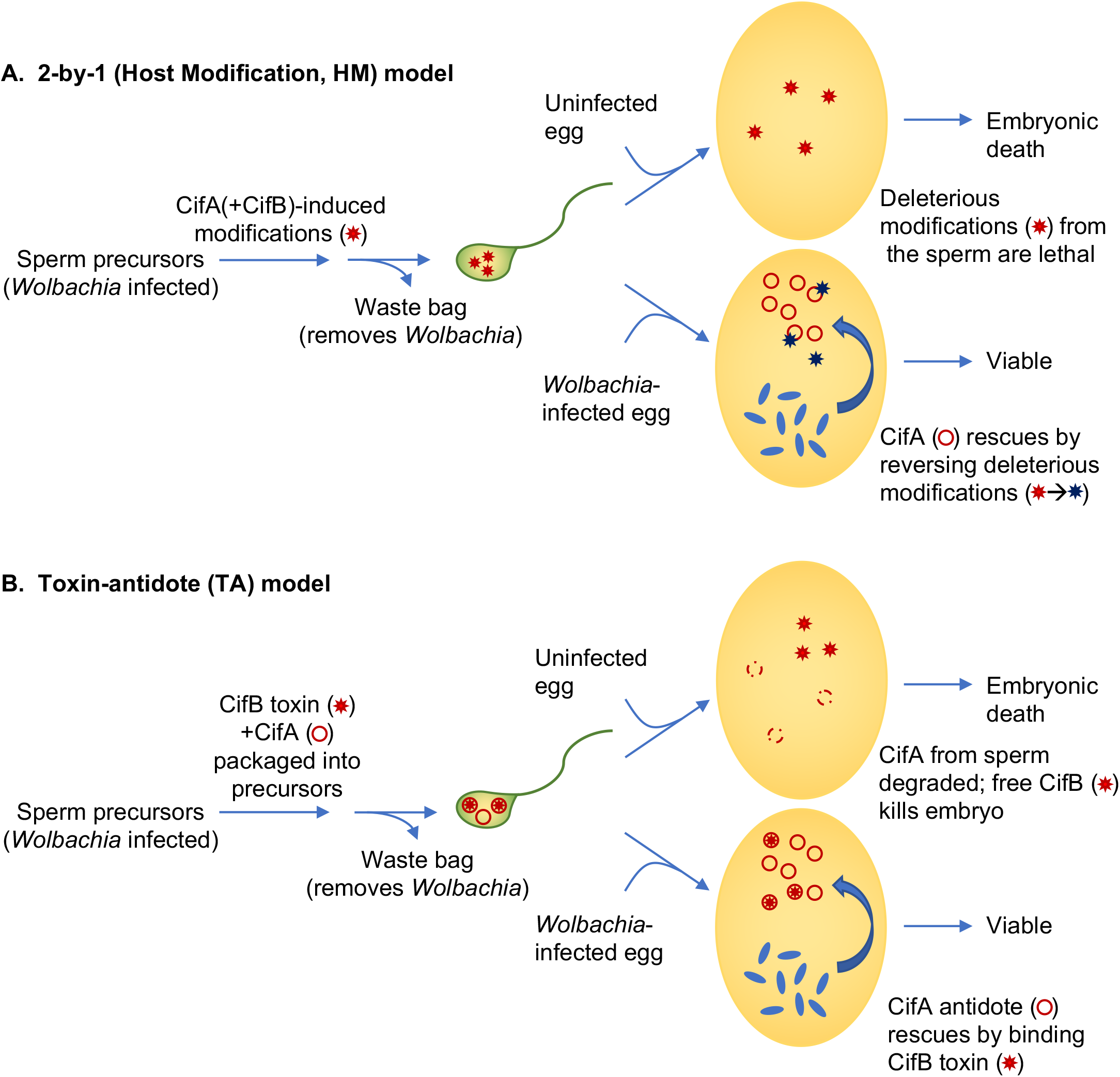
Models for cytoplasmic incompatibility (CI) caused by *Wolbachia*. A. The 2-by-1 host modification (HM) model. Although the 2-by-1 model is strictly a genetic description, the most recently articulated version has been interpreted in the context of the HM mechanistic model shown here (19). CifA is regarded as the key CI-inducing factor that is responsible for modifications of sperm; CifB in this scheme has an undefined accessory role within the testes that is required for CI induction. In the fertilized egg, if CifA is secreted by infecting *Wolbachia* (blue ovals), it works in opposite fashion to reverse the sperm modifications and rescues embryonic viability. *Wolbachia* are known not to be incorporated into mature sperm and are instead eliminated along with other organelles in the “waste bag” during spermiogenesis. There are multiple additional variants of the HM model that predate identification of the Cif proteins. B. The toxin-antidote (TA) model. In this model, CifB is the fundamental CI-inducing factor and is postulated to be imported into the mature sperm (along with CifA); CifA may promote CifB levels or packaging into sperm or protect sperm precursors from CifB-induced toxicity. Upon fertilization, or possibly before, CifB enzymatic activity –either from the CidB deubiquitylase or CinB nuclease– alters the sperm-derived chromosomes in some way. CifB is proposed to be relatively long-lived and CifA, the antidote that binds CifB, short-lived, so additional high-level expression of CifA in the egg is required to counter CifB activity.

**Figure 2.**
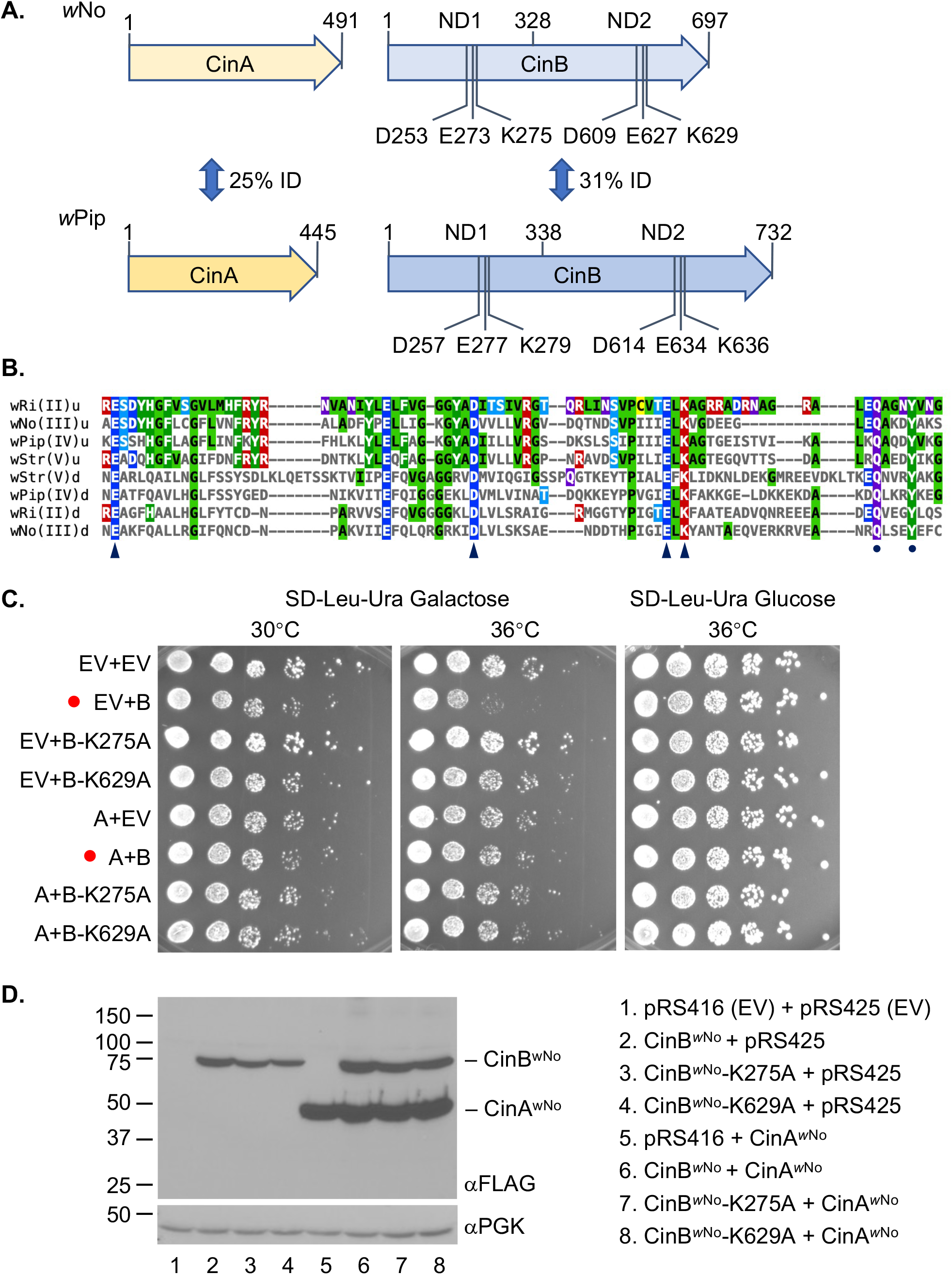
The *w*No-derived CinA and CinB proteins. A. Comparison of the *w*No and *w*Pip CinA and CinB CI factors. The *w*No proteins are from the type III clade of CI factors; the *w*Pip proteins belong to the type IV branch. Five base pairs (bp) separate the stop codon of *w*No *cinA* from the start codon of *cinB;* in the *w*Pip *cinA-cinB* operon, this gap is 51 bp. All CinB proteins include two intact nuclease domains (NDs), which both appear to be necessary for DNase activity and biological function. ID, identity. B. Core sequences of the PDDEXK nuclease domains (NDs) from the four known clades of CifB factors thought to encode active nucleases (types II-V). The conserved residues that constitute the active site based on the CinBo^*w*Pip^ (type IV) crystal structures (23) are indicated by arrowheads. A conserved QxxxY motif just downstream of the active site residues is marked by dots; this motif is characteristic of RecB-like nucleases and has been suggested to function in DNA binding (32). Alignments were done with Clustal Omega, and the figure was made using MView (1.63). u, upstream or N-terminal ND; d, downstream or C-terminal ND. C. Growth assays of yeast expressing *w*No *cinA* and *cinB* alleles. BY4741 cells were transformed with plasmids expressing the indicated alleles from a galactose inducible *GAL1* promoter, and cultures were spotted in serial dilution onto selection plates with the indicated carbon source and grown for 2.5 days at either 30°C or 36°C. Red dots highlight strains expressing WT CinB^*w*No^ with and without CinA^*w*No^ co-expression. EV, empty vector; A, CinA^*w*No^; B, CinB^*w*No^. D. Expression levels of different CinA and CinB proteins in the same BY4741 transformants from panel C were measured by immunoblot analysis. Both proteins were tagged with a FLAG epitope. The anti-PGK immunoblot served as a loading control.

CidA^*w*Pip^ and CinA^*w*Pip^ bind the cognate CidB^*w*Pip^ and CinB^*w*Pip^ proteins with high affinity, but the binding does not block the enzyme active sites of the latter proteins (10, 12). Nevertheless, coexpression in yeast of the cognate CifA protein from each operon suppresses the toxicity observed if CidB or CinB is expressed by itself (10). These observations suggest that suppression results from interference of the CifA factors with CifB enzyme-substrate interactions or from enzyme relocalization within the cell. Expression of the CifA proteins by themselves is not deleterious to yeast (17).

Results from yeast growth studies strongly parallel effects seen by transgenic germline expression of the same Cif proteins in *Drosophila melanogaster*. For example, female germline expression of CidA^*w*Mel^ or CinA^*w*Pip^ was sufficient for rescue of transgenic CI caused by expression of the cognate *cifA-cifB* genes in the male germline (12, 18). The enzymatic activity of the CifB enzymes is required for full CI in transgenic flies, as is true for yeast toxicity (10, 12, 19). Moreover, orthologous α-karyopherin proteins were shown to suppress CI or growth inhibition in both flies and yeast (17).

By contrast, induction of CI in flies required not only expression of CifB, as would be predicted from the yeast studies, but also that of the partner CifA (12, 14). This requirement for both *cifA* and *cifB* for transgenic CI induction and *cifA* alone for rescue has been termed the ‘2-by-1’ model (18). The dual gene requirement for induction has called into question the reliability of yeast growth assays as a surrogate for CI analysis in transgenic insects even though the CI rescue activity of CifA proteins was initially predicted from yeast studies (10).

In the current study, we analyze a divergent *cinA-cinB* system from the *w*No *Wolbachia* strain, which naturally infects *D. simulans* (20). We had shown previously that *w*No-infected *D. simulans* display embryonic cytological defects in incompatible crosses that are similar to those observed with incompatible transgenic *D. melanogaster* crosses involving *cidA-cidB* or *cinA-cinB* from *w*Pip (12). However, we had not tested whether transgenic *cinA*-*cinB^w^*^No^ behaves similarly. Here we find that this is indeed the case. Unexpectedly, however, male expression of *cinB^w^*^No^ alone induced strong CI in transgenic flies, and this was fully rescued by cognate *cinA^w^*^No^ expression in the female germline. Parallel results were seen with *cinA-cinB*^*w*No^ expression in yeast where it was further shown that the nuclease active site must be intact to observe toxicity. These findings indicate that catalytically active CifB is the fundamental CI-inducing factor and that CifA is the specific rescue factor. Our results also bear on current models for CI mechanisms. In particular, they argue against the generality of the 2-by-1 model and models that posit a primary role for CifA in CI induction (**Figure 1**).

## Results

### Sequence comparisons of CinA and CinB homologs

As noted above, we had previously shown that the CinA and CinB factors from *w*Pip *Wolbachia* are sufficient to recapitulate key features of CI in transgenic *D. melanogaster* (12). Primary sequence comparisons divide the *cifA-cifB* gene pairs from different *Wolbachia* strains into five distinct clades or types (21, 22). In four of these, types II-V, the downstream gene is predicted to encode an active nuclease. The CinB^*w*Pip^ nuclease is a type IV Cif protein. Because *w*Pip(Pel) encodes both *cidA-cidB* (type I *cif*) and *cinA*-*cinB* gene pairs, one cannot infer which of the two, if not both, is responsible for causing CI. We therefore had turned to an analysis of *D. simulans* infected with *w*No, a *Wolbachia* strain with only a single *cinA-cinB* operon, to verify that the embryonic cytological features of CI associated with a nuclease-encoding operon were similar to those seen with *Wolbachia* strains such as *w*Mel, which only have a *cidA-cidB* (DUB-encoding) locus (12). However, in our previous analysis of *w*No-induced CI, we had not tested whether the CinA and CinB factors from *w*No could recapitulate CI transgenically.

The *w*No CinA and CinB proteins have diverged considerably from the homologous proteins from *w*Pip (**Figure 2A**). Nevertheless, the nuclease catalytic residues are conserved in CinB^*w*No^, and it is also predicted to have two active PD-(D/E)XK (PDDEXK) nuclease domains (**Figure 2A, B**). In the N-terminal nuclease domain (NTND) of CinB^*w*Pip^, the catalytic residues coordinate a divalent cation, whereas no cation binding was observed in the C-terminal ND (CTND) (23). Despite this, mutating the catalytic residues in either domain abolishes DNase activity and blocks CI-like induction (12). The conservation of these residues in both domains among CifB types II-V (**Figure 2B**) predicts that CinB^*w*No^ will also function in CI in a way that depends on its nuclease activity.

### Analysis of CinA^wNo^ and CinB^wNo^ expression in budding yeast

Growth analysis of *S. cerevisiae* expressing different *Wolbachia* Cif proteins has generally proven to be a reliable surrogate for transgenic CI analysis in insects (10, 12, 15). Expression of CidB proteins from *w*Pip or *w*Ha was found to cause temperature-sensitive growth defects in yeast that were suppressed by co-expression of their cognate CidA proteins (10, 17). The same was true for CinA^*w*Pip^ and CinB^*w*Pip^ expression (10). We therefore transformed yeast with plasmids carrying *cinA*^*w*No^, *cinB*^*w*No^, or both under galactose-inducible promoters and examined growth of the transformants (**Figure 2C**). Galactose-induced production of CinB^*w*No^ caused a reproducible defect in growth at higher temperatures, although the impairment was less severe than that previously seen with CinB^*w*Pip^. Importantly, the growth deficiency was relieved by CinA^*w*No^ co-expression.

When a predicted catalytic lysine in either the NTND or CTND (K275 and K629, respectively) was mutated to alanine, yeast growth impairment was abrogated (**Figure 2C**; **Suppl. Figure S1**). Unlike what had been observed with CinB^*w*Pip^, the active-site mutations caused a partial reduction in levels of the protein. This effect was relatively mild in the BY4741 strain background (**Figure 2D**) but was more marked in a W303 strain (**Suppl. Figure S1**). The modest reduction of CinB^*w*No^ mutant protein amounts seen in the former strain would not predict complete loss of growth inhibition. Interestingly, co-expression of CinA^*w*No^ reproducibly increased the levels of the cognate CinB^*w*No^ proteins, whether WT or mutant. This suggests that CinB^*w*No^ might be susceptible to degradation in yeast and that the (predicted) binding to CinA^*w*No^ may protect it from such turnover.

In summary, these data indicate that the CinA^*w*No^-CinB^*w*No^ cognate pair behaves similarly to those encoded by other *Wolbachia cifA-cifB* genes previously analyzed in yeast.

### Cognate-specific binding preferences of *w*No and *w*Pip CinA-CinB pairs

The ability of CinA^*w*No^ to suppress the growth defect of yeast expressing CinB^*w*No^ (and to elevate CinB^*w*No^ levels) suggested that the two proteins interact, as was known to be true for CinA^*w*Pip^-CinB^*w*Pip^, which form a tight complex (Kd, 25 nM) (12). To test this directly, recombinant His6-tagged CinA and glutathione-S-transferase (GST)-tagged CinB proteins derived from *w*No and *w*Pip were co-expressed in *Escherichia coli* and tested for binding using GST pulldown assays (**Figure 3A**). As expected, CinA^*w*Pip^-CinB^*w*Pip^ showed a strong interaction (**Figure 3A**, lane 1), which was likely a 1:1 complex based on our recent crystallographic study (23). Similarly, CinA^*w*No^-CinB^*w*No^ displayed a readily detectable interaction (lane 4). By contrast, the noncognate pairs showed only weak cross-binding (lanes 2, 3). Similar results were found when lysates from separate *E. coli* transformants expressing either recombinant CinA^*w*No^ or CinB^*w*No^ were mixed and evaluated by GST-pulldown analysis (Suppl. Figure S2). Consistent with these binding data, suppression of CinB-induced yeast growth defects by CinA proteins showed cognate specificity: rescue was only seen when the CinA and CinB proteins came from the same *Wolbachia* strain (**Figure 3B**).

**Figure 3.**
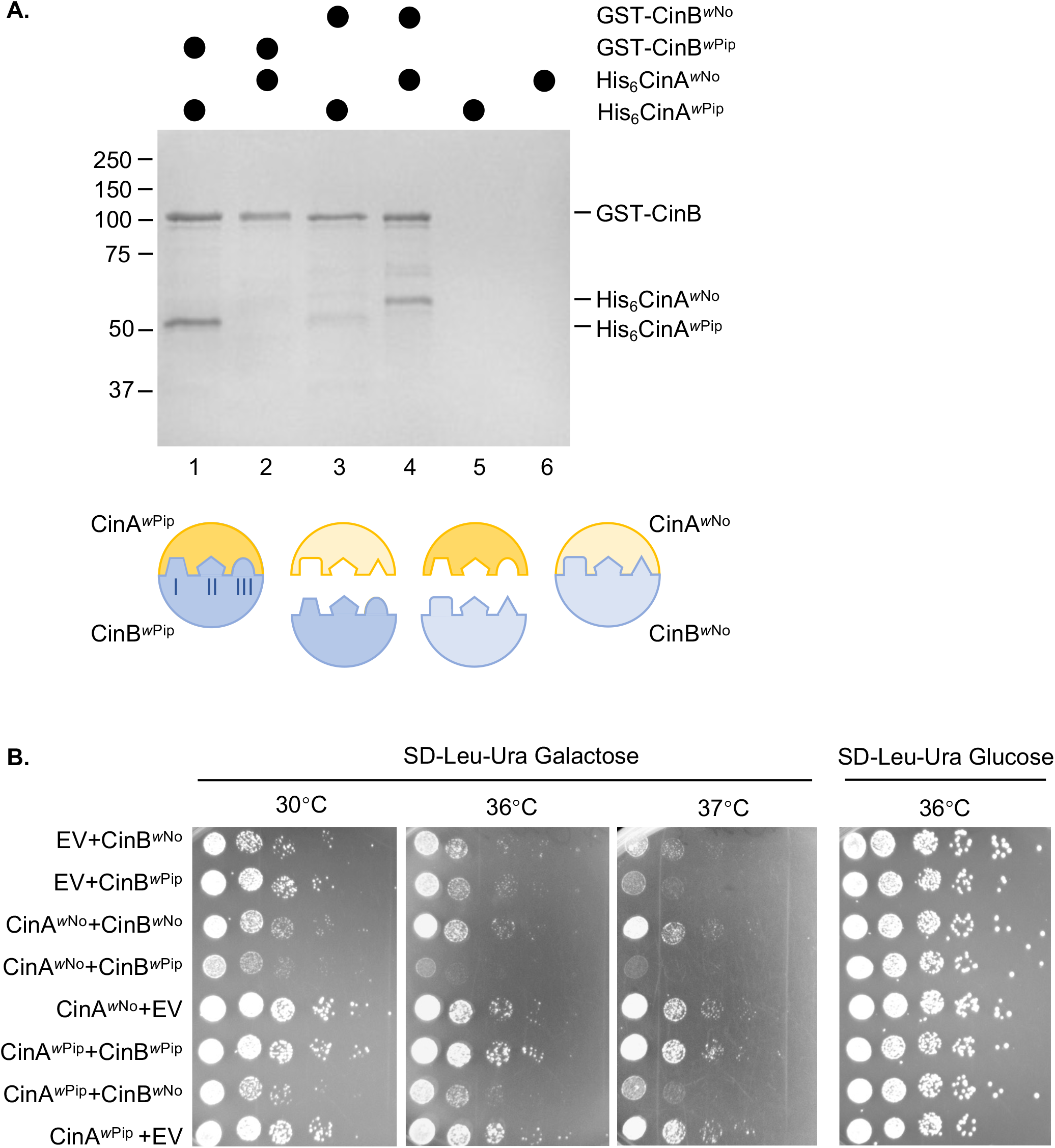
*w*No CinA and CinB form a cognate-specific protein complex. A. GST-pulldown analyses were done with recombinant proteins co-expressed in *E. coli*. Lysates containing the indicated proteins were bound to a glutathione resin. After washing, proteins were eluted with reduced glutathione, resolved by SDS-PAGE, and visualized with GelCode Blue. Cognate protein pairs are in lanes 1 and 4. Size standards, in kDa, are indicated at left. The cartoon interpretation below illustrates the complementary tripartite interfaces between cognate CinA-CinB pairs (23). B. CinA-CinB from *w*No and *w*Pip show cognate specificity in yeast growth rescue. Growth assays were done with W303 yeast expressing *cinA* and *cinB* alleles from *w*No or *w*Pip. Cultures were spotted in serial dilution onto selection plates and grown for 2.5 days at either 30°C, 36°C, or 37°C. EV, empty vector (p425GAL1 for *cinA* genes, pRS416GAL1 for *cinB*). As noted previously (17), CinA^*w*No^ enhances toxicity when expressed with noncognate CinB proteins.

### CinB^*w*No^ is sufficient for transgenic CI induction and CinA^*w*No^ is sufficient for its rescue

There are still only a handful of examples where the ability of specific *cifA-cifB* gene pairs to recapitulate CI by transgenic germline expression in insects has been demonstrated. Moreover, it has been suggested that there may be additional *Wolbachia* genes that are responsible for CI (24). We therefore tested whether the highly divergent *w*No-encoded CinA and CinB proteins (**Figure 2A**) were indeed capable of causing CI.

In contrast to yeast growth impairment, expression of both *cifA* and *cifB* genes in the *D. melanogaster* male germline has been reported to be necessary for triggering transgenic CI (18). Both proteins are expected to be present in mature sperm, although only CidA^*w*Pip^ has been directly detected there (25). This has raised questions about whether CifA or CifB is the primary inducer of CI (**Figure 1**) (19).

We first created transgenic flies that expressed the *cinA^w^*^No^-*cinB^w^*^No^ coding sequences linked by a picornavirus T2A sequence; the latter element is expected to lead to expression of the two proteins in roughly equal amounts from a single mRNA (26). Expression of the CinA-T2A-CinB (*w*No) construct in males that were crossed to wild type (WT) females caused a nearly complete loss of viable embryos based on egg hatch rates (**Figure 4A**). The *cinA*^*w*No^ gene by itself caused no reduction in viability, as expected, despite high relative mRNA levels (Suppl. Figure S3).

**Figure 4.**
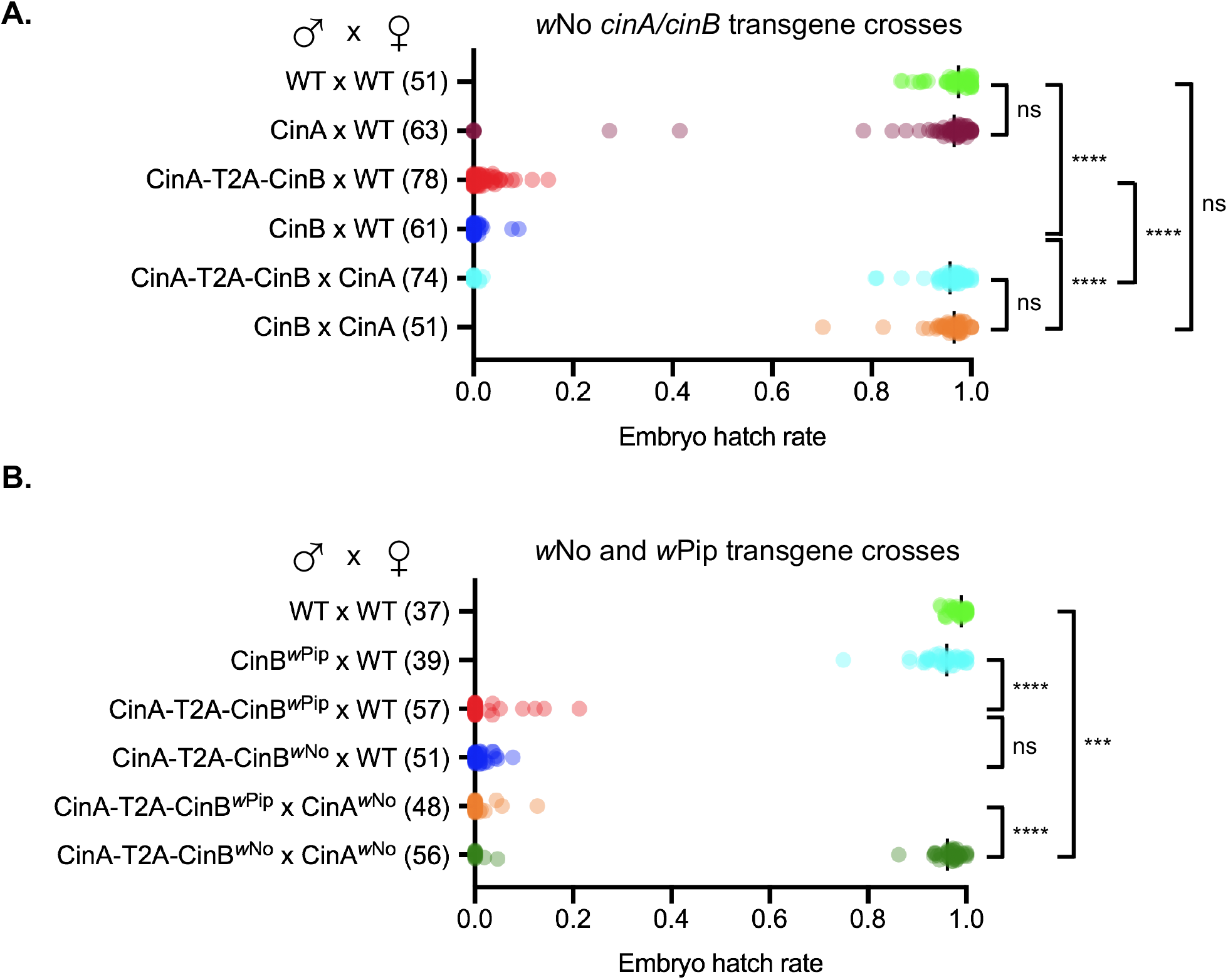
Paternally derived CinB^*w*No^ by itself causes strong embryonic lethality. A. Male germline expression of a *cinB*^*w*No^ transgene causes post-zygotic lethality and is fully rescued by *cinA*^*w*No^ transgene expression in females based on egg hatch-rate analysis. The number of one-on-one matings used for each cross is shown in parentheses. B. The *cinA*^*w*No^ transgene does not rescue noncognate *cinA-cinB*^*w*PiP^-induced embryonic lethality. All the strains employed for the test crosses contained the *MTD-GAL4* driver except for the CinA-T2A-CinB^*w*No^ strain. The *cinA* and *cinB* coding sequences in the latter construct were linked by a T2A viral sequence that causes the ribosome to terminate and then reinitiate translation within the linker, expressing CinA and CinB as separate proteins. We note that the control rescue cross of *cinA-cinB*^*w*Pip^ with *cinA*^*w*Pip^ (female) failed for unknown reasons. ***P < 0.001, ****P < 0.0001 by ANOVA with multiple comparisons between all groups.

Surprisingly, expression of the *cinB*^*w*No^ gene alone in the male germline induced a severe reduction in viable embryos that was indistinguishable from that induced by the *cinA*^*w*No^-*cinB*^*w*No^ pair. This embryonic lethality was completely reversed by expression of *cinA*^*w*No^ in the female germline (**Figure 4A**). We also repeated our earlier analysis of transgenic CI in *D. melanogaster* caused by the *cinA*^*w*Pip^-*cinB*^*w*Pip^ pair (12) (**Figure 4B**). As reported previously, both *cinA*^*w*Pip^ and *cinB*^*w*Pip^ were needed for inducing strong embryonic lethality, in contrast to the sufficiency seen with *cinB*^*w*No^. Expression of *cinA*^*w*No^ in the female germline did not rescue viability in crosses to males with the heterologous *cinA*-T2A-*cinB*^*w*Pip^ construct. In summary, these data strongly suggest that CI, at least in this transgenic model, does not require the *cifA* component of the *cifA-cifB* locus, in contrast to previous suggestions.

Although rescue of embryo viability by *cinA*^*w*No^ was robust and strongly implicated CI as the cause of the observed male sterility, it was important to document that the cytological features of transgenic CI induced by *cinB*^*w*No^ resembled natural CI rather than some other form of male sterility. Typically, CI leads to arrest during the first zygotic mitotic division due to asynchronous chromosome condensation and separation in the juxtaposed male and female pronuclei (4, 27). Not all embryos will necessarily arrest at this stage, but nuclear division stalls in a large fraction of embryos prior to the blastoderm stage (10, 11, 16).

We analyzed embryo cytology 1-2 hours after egg deposition in transgenic CI crosses (**Figure 5A-E**). Either expression of CinB^*w*No^ alone or in combination with CinA^*w*No^ in the male germline caused most resulting embryos to arrest early in development. Expression of CinA^*w*No^ in the female germline strongly countered the arrest phenotype in both cases. When embryos from CinB^*w*No^ (male) x WT crosses were examined after a shorter (30 min) egg-laying period, we were able to observe first zygotic division defects; these ranged from asynchronous condensation of chromosomes between the juxtaposed male and female pronuclei to chromatin bridging in late anaphase or telophase (**Figure 5F**). Therefore, the male sterility which occurs because of transgenic CinB^*w*No^ expression in males bears the hallmarks of natural CI previously observed in *w*No-infected *D. simulans* and other examples of CI (12).

**Figure 5.**
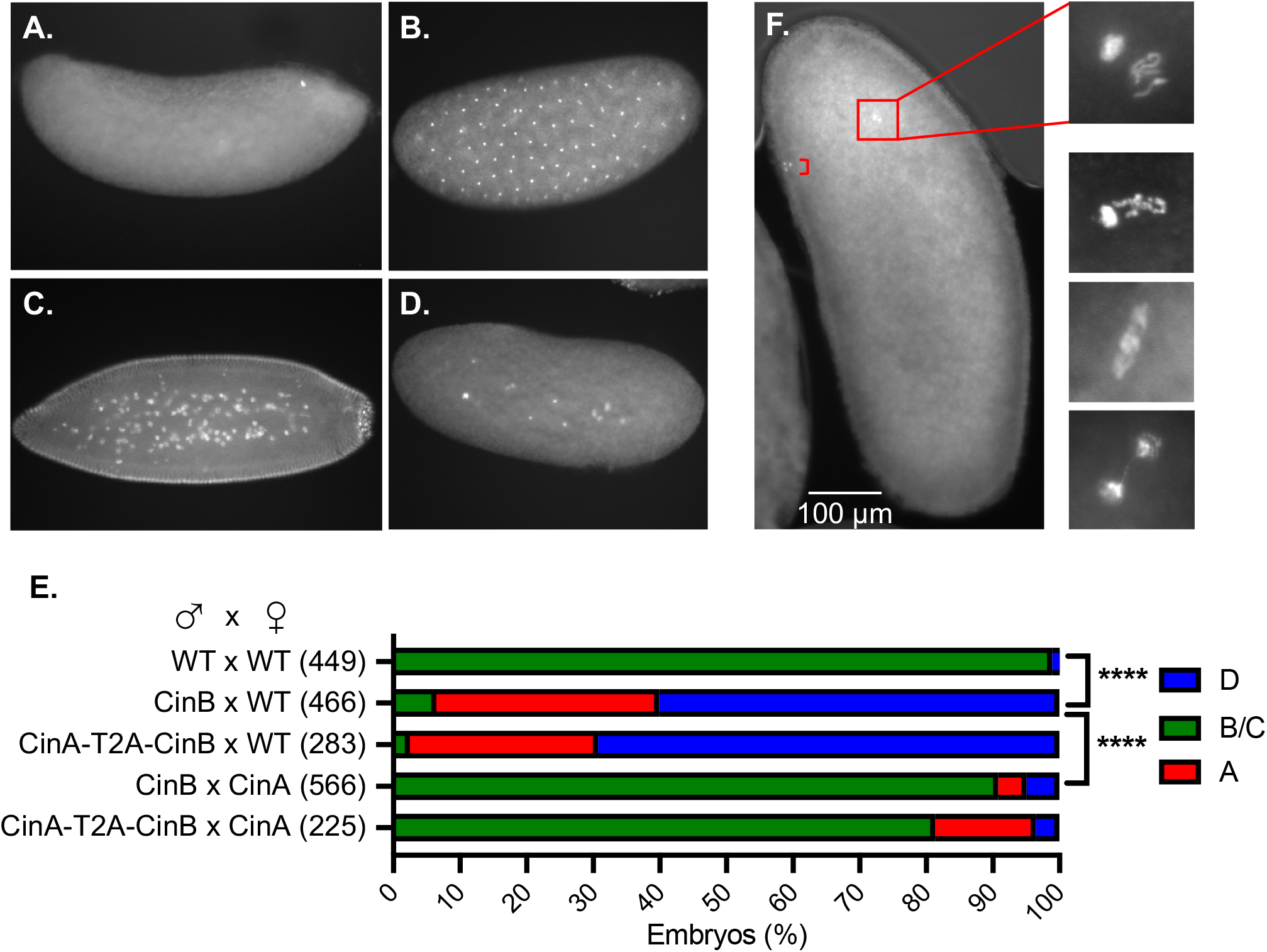
CI-like embryonic defects caused by expression of CinB^*w*No^ or CinA-CinB^*w*No^ in males. A–D. Representative images of early embryos with DNA staining by propidium iodide. (A) Unfertilized or very early-arrest embryo; (B) normal embryo at ~1 h of development; (C) normal embryo at ~2 h of development; and (D) embryo with early mitotic failure. E. Quantification of embryo cytology into classes A-D. Embryos exhibiting normal cytology at 1 to 2 h were grouped together and are shown in green. ****, P< 0.0001 by χ2 test comparing normal (B and C) and abnormal (A and D) cytological phenotypes. All transgenic strains were confirmed by PCR (12), and all strains used in the test crosses carried the *MTD-GAL4* driver except for the CinA-T2A-CinBo’’^No^ strain. The number of embryos examined for each cross is in parentheses. F. Images of Hoechst 33342-stained embryos from transgenic CI crosses between transgenic CinB^*w*No^ males and (uninfected) wild-type females showing apposed but asynchronous male and female pronuclei with defects at the first zygotic mitosis. Embryos were fixed after allowing 30 min for egg laying. Bracket in the low magnification image marks the three polar bodies.

## Discussion

The current study includes several findings relevant to current models for the mechanism of *Wolbachia-mediated* CI. The CinA and CinB CI factors from *w*No behave in a manner very similar to the previously analyzed Cin (and Cid) proteins of *w*Pip. Despite their strong sequence divergence, the CinB nucleases from both *w*No and *w*Pip can induce CI in fruit flies and toxicity in yeast. CinA^*w*No^ is sufficient for rescue of CI in transgenic *Drosophila* and suppression of toxicity in yeast. Paralleling their growth effects, the *w*No CinA and CinB proteins bind one another in a cognate-specific fashion. Moreover, mutation of predicted catalytic residues of the nuclease active centers in CinB^*w*No^ prevents its toxicity when expressed in yeast. Most importantly, expression of CinB^*w*No^ alone in the male germline of transgenic flies induces highly penetrant CI. This embryonic lethality can be fully rescued by CinA^*w*No^ expression in females.

One significant discrepancy in previous studies of the *cif* genes was the observation that yeast growth defects caused by heterologous Cif protein expression and transgenic male sterility caused by these same factors showed different *cif* gene dependencies. Specifically, *cifB*-encoded proteins were sufficient for growth impairment in yeast, whereas expression of both *cifA* and *cifB* genes in the *D. melanogaster* male germline was needed to trigger transgenic CI (12, 18). On the other hand, the ability of cognate *cifA* genes by themselves to rescue transgenic CI matched what had been seen by yeast suppression analysis (10, 12).

The requirement for both CifA and CifB in transgenic CI induction but only CifA for rescue has been enumerated as the ‘2-by-1’ model (**Figure 1A**) (18). Although this genetic summary is usually interpreted in the context of a host modification (HM) mechanism for CI, it can be accommodated by a toxin-antidote (TA) scheme as well (**Figure 1**). However, recent versions of the 2-by-1 model (for example, (19)) have posited a primary role for CifA in CI induction in which CifA, by an unknown mechanism, causes modifications in sperm precursors that, following fertilization, will be lethal to the embryo if not reversed, also by CifA, in the egg. CifB, in this view, has only an ancillary role in CI induction, possibly by maintaining proper levels or activity of CifA during spermatogenesis. While the plausibility of this model could be questioned on several grounds, it does make a straightforward prediction, which is that CifA is strictly necessary for CI induction.

Our data demonstrate that the *w*No CifA (CinA) protein is dispensable for CI induction and instead show that *w*No CifB (CinB) is sufficient; this embryonic lethality is completely blocked following mating with females expressing CinA^*w*No^ in the germline (**Figures 4 and 5**). These findings align well with results on yeast toxicity caused by CinB^*w*No^ alone and its suppression by CinA^*w*No^ (**Figures 2C, 3B**). In a recent preprint, expression of *w*Pip CidA and CidB proteins in the malaria vector *Anopheles gambiae* has been reported (28). Expression of CidB^*w*Pip^ by itself in male mosquitoes induced very strong transgenic CI, and CidA^*w*Pip^ in females was sufficient to block the embryonic lethality caused by crossing to transgenic *cidB*^*w*Pip^ male mosquitoes. These and our results together therefore argue that the fundamental inducers of CI are the CifB proteins, and this is true for both CinB nuclease and CidB deubiquitylase *Wolbachia* CI factors. Three different model organisms −*D. melanogaster, A. gambiae*, and *S. cerevisiae*–used for transgenesis studies of the *cif* genes have now yielded evidence for a ‘1-by-1’ genetic scheme in which CifB is the lone modification factor or toxin and CifA is the rescue factor or antidote.

Nevertheless, both the A and B factors from other *Wolbachia cif* operons were previously shown to be necessary for transgenic CI induction in *Drosophila* (11, 12), and here we have reproduced our original finding showing the *cinB*^*w*Pip^ is not sufficient for inducing strong transgenic CI but requires *cinA*^*w*Pip^ co-expression (**Figure 4B**). The exact reason for the different CI gene requirements using genes from the same CI operon for expression in different insect models (*Drosophila* versus *Anopheles*) or from CI operons of different *Wolbachia* strains is not known, but it likely derives from the level and localization of transgenic expression of these proteins in the male germline (see (18) for fuller discussion). The co-expression data in yeast suggest that CifA might stabilize the CifB protein, although this remains to be tested (**Figure 2D**). The baker’s yeast model has been a useful predictor of the cellular effects of the CI factors even though yeasts are not known to be natural hosts to *Wolbachia*.

By themselves, the current data do not distinguish between HM and TA mechanisms. The TA mechanism, but not HM, explicitly requires binding between modifier/toxin and rescue/antidote factors (**Figure 1**). Given the results in the current work, this binding should occur between CifB brought in by sperm and CifA expressed in the female germline. Support for this has recently been garnered using high-resolution crystal structures for several CifA-CifB binary complexes (23). From these structures, it was found that the interfaces between the A and B factors are dominated by charged and polar residues. Amino acids in these interfaces could be mutated to eliminate binding; this blocked the ability of the CifA rescue factors to suppress cognate CifB toxicity in yeast and to rescue transgenic CI in flies. If complementary mutations were made at the interfaces to partially restore CifA-CifB binding, yeast toxicity caused by the mutant CifB was suppressed by co-expression of the CifA interface mutant.

The current work aligns results from different model systems and makes clear which CI factors are fundamentally responsible for the induction and rescue of CI. Ultimately, determining the mechanism of CI will require identifying the relevant molecular targets of the CifB enzymes in the host and the timing of their action, and determining how direct binding by the CifA factors blocks or reverses CifB action *in vivo* (17). Neither *in vitro* CinB nuclease activity nor CidB deubiquitylase activity is diminished by cognate CifA binding (10, 12). Therefore, CifA proteins may block or reverse enzyme-substrate interactions or alter enzyme localization (or both) *in vivo*. Whatever the CI mechanisms, they are likely to impact conserved physiological pathways given the wide host range of *Wolbachia* that cause CI and the even broader range of species in which heterologous expression of the CI factors mimics a toxin-antidote system. Understanding these mechanisms will allow more rational manipulation of CI for applications in crop protection and disease vector control.

## Methods

### Yeast and bacterial strains and plasmid construction

MHY1774 (W303-1A) and MHY10139 (BY4741) yeast strains were used for yeast growth assays. *E. coli* Top10F’ cells were used for DNA cloning, and Rosetta BL21(DE3 pLysS) cells were employed for protein expression. PCR products were generated with the primers listed in Supplementary Table 2 using HF Phusion DNA polymerase (New England Biolabs).

For galactose-inducible expression of proteins in yeast, pRS425GAL1 (*LEU2*) and pRS416GAL1 (*URA3*) expression plasmids were used (29). Constructs p425GAL1-Flag-CinA^*w*No^ and p416GAL1-Flag-CinB^*w*No^ were described earlier as were the *w*Pip equivalents (17). Mutations were introduced into Flag-CinB^*w*No^ in p416GAL1 by Quikchange (Agilent; mutagenesis using primers listed in Supplementary Table 2).

For protein expression in *E. coli* and transgene expression in *D. melanogaster*, coding sequences of CinA^*w*No^, CinB^*w*No^ and CinA^*w*No^-T2A-CinB^*w*No^, optimized for expression in *Drosophila*, were synthesized by Genscript. To make transgenic flies, the pTiger vector, bearing a Gal4-binding UAS site was used. CinA^*w*No^, CinB^*w*No^ and CinA^*w*No^-T2A-CinB^*w*No^ coding sequences were separately cloned into pTiger using KpnI and SpeI restriction sites. For expression of proteins in *E. coli*, pET28a-pp, which encodes an N-terminal His tag, and pGEX-6P-1, which encodes an N-terminal GST tag, were used. The *cinA*^*w*No^ gene was subcloned from pRS425-CinA^*w*No^ into pET28a-pp using BamHI/SalI sites; *cinB*^*w*No^ was subcloned from pRS416-CinB^*w*No^ into pGEX-6P-1 using BamHI/NotI sites. pGEX-6P-1-CinB^*w*Pip^ was reported previously (12), while the coding sequence for CinA^*w*Pip^ was subcloned from pUAS-attb-CinA^*w*Pip^ (12) into pET28a-pp NdeI/XhoI sites. Also see Supplementary Table 1.

### Yeast growth analysis

MHY1774 or MHY10139 yeast were transformed with the plasmids indicated in the figures. To test growth, cultures were grown at 30°C in SD raffinose media lacking uracil and leucine for 2 days before spotted in six-fold serial dilution onto SD plates lacking uracil and leucine and containing either 2% galactose or glucose.

For immunoblot testing of *Wolbachia* protein expression in yeast, co-expression cultures in SD-raffinose lacking uracil and leucine were diluted to 0.2 OD600 in SD-galactose medium lacking uracil and leucine and cultured for 12-16 h at 30°C until reaching an OD600 of 0.8-1.0. Cells were harvested and treated for 5 min at room temperature with 0.1 M NaOH (30). Cell pellets were lysed in SDS sample buffer and clarified lysates were analyzed by immunoblotting (17). Mouse anti-FLAG M2 (Sigma, 1:10,000) or mouse anti-PGK (yeast phosphoglycerate kinase; Abcam, 1:10,000) were used for immunoblotting along with sheep anti-mouse-HRP NXA931V secondary antibody (GE Healthcare, 1:5,000 or 1:10,000, respectively). All yeast growth and Western blot data shown are representative of at least two biological replicates.

### Protein-binding analysis

To test interactions between various CinA and CinB proteins, GST fusion protein pull-down assays were performed. Two different expression methods were used. In the first, pET-28a-pp-CinA (conferring kanamycin resistance) and pGEX-6P-1-CinB (conferring ampicillin resistance) plasmids were co-transformed into Rosetta *E. coli* BL21(DE3 pLysS). Bacteria were grown at 37°C in 75 mL LB (Luria Broth) medium containing kanamycin and ampicillin to OD_600_=0.6-0.8, induced with 600 μM IPTG (isopropyl-β-thiogalactoside) and incubated ~20 h at 18°C. Cultures were centrifuged at 4°C, 7,000 × *g* for 10 min.

Cell pellets were resuspended in GST-tag lysis buffer (50 mM Tris-HCl pH 7.5, 150 mM NaCl, 0.01% Tween-20, 10 mM imidazole pH 8.0, 2 mM phenylmethylsulfonyl fluoride (PMSF), 20 μg/mL DNase I and 1 mg/ml chicken egg-white lysozyme), kept on rotator at 4°C for 30 min and then lysed by sonication. Crude lysates were clarified by centrifuging at 21,000 x *g* for 30 min at 4°C. Clarified lysates were incubated with 50 μL Pierce glutathione-agarose beads (Thermo, USA) pre-washed with wash buffer (50 mM Tris-HCl pH 8.0, 150 mM NaCl) and rotated at 4°C for 1 h. Beads were washed six times with 1 mL wash buffer, and proteins were eluted by adding two bed volumes of elution buffer (50 mM Tris-HCl pH 8, 150 mM NaCl, 20 mM reduced glutathione) and incubated on a rotator at room temperature for 15 min. Inputs and eluted fractions were resolved by SDS-PAGE. The proteins were visualized by GelCode Blue staining (ThermoFisher, USA) and imaged on a G:Box (Syngene).

In the second method, the expression plasmids were transformed individually into competent Rosetta cells. Cultures were grown and pelleted as above. Cells expressing His6-CinA were resuspended in His-tag lysis buffer (50 mM Tris-HCl pH 7.5, 150 mM NaCl, 0.01% Tween-20, 10 mM imidazole pH 8.0, 2 mM PMSF, 20 μg/mL DNase I, and 1 mg/ml lysozyme), while those expressing GST-CinB were resuspended in GST-tag lysis buffer. Cells were incubated on a rotator for 30 min at 4°C, then lysed by sonication as above. Clarified GST-CinB-containing lysates were first incubated with glutathione-agarose resin and rotated at 4°C for 1 h. After removing the flow-through, the beads were washed three times with 1 mL GST-tag wash buffer, followed by addition of clarified His6-CinA lysates and rotation at 4°C for 1 h. The beads were washed five times with 1 mL wash buffer, with all subsequent steps as in the previous method.

### Hatch-rate analysis of transgenic D. melanogaster crosses

To generate the *w*No Cif transgenic flies used for hatch-rate analyses, DNA constructs were sent to BestGene, Inc. for microinjection into *D. melanogaster* embryos. Flies transgenic for *cinA^w^*^Pip^, cinB^*w*Pip^ and *cinA-T2A-cinB*^*w*Pip^ were generated previously (12). White Canton-S (wCS; WT; #189), MTD driver (#31777, GAL4) fly strains were gifts or were obtained from the Bloomington Stock Center. Fly strain #9744 (containing an attP insertion site on the third chromosome) was used for all gene constructs for site-directed attP/B integration by the PhiC31 integrase except the #9723 strain was chosen for site-specific integration of *cinA*^*w*No^ on the second chromosome. Flies were reared on standard corn meal-based solid media.

All flies for the hatch-rate assays were homozygous for the integrated genes except for the *cinA*-T2A-*cinB*^*w*No^ line, which showed a high degree of sterility when virgin red-eyed individuals were mated. This made it unfeasible to maintain a homozygous line; the sterility was likely due to leaky transgene expression, which is known to occur with the UAS system (10). For *cinA*-T2A-*cinB*^*w*No^, red-eyed progeny of white-eyed males and red-eyed virgin females were used for maintenance of the heterozygous line as well as for hatch-rate experiments.

Except for the *cinA*-T2A-*cinB*^*w*No^ flies, parental flies used in the hatch-rate experiments were generated by crossing *MTD-GAL4* virgin females to the corresponding transgenic males. The crosses were maintained at ~22°C (temperature was temporarily lowered to 18°C for overnight virgin collection) on a standard diet. Virgin flies of the desired genotype and sex were collected, aged at 25°C for 2-4 additional days, and used to set up 1×1 crosses for hatch-rate determination. For each cross, a virgin female and male were mated as described (12). All crosses were incubated at 25°C overnight (~17 h) (initial incubation), and the original apple juice plates were set aside and replaced with freshly yeasted plates, which were kept at 25°C for 24 h (additional incubation). Adult flies were then removed and frozen at −80°C for future expression analysis. Both sets of plates were incubated at 25°C for another 24 h before the number of hatched and unhatched embryos was counted. Embryo numbers from the two sets of plates were pooled, and any mating pair with fewer than 15 total embryos laid were discarded. The counting was not blinded. One-way ANOVA with multiple comparison statistical analysis was performed using GraphPad Prism (v. 9) software.

### Cytological analysis of embryos

To prepare embryos for cytological analysis, ~100 males and ~100 females (2-4-day old virgins) were used. The methods were described previously (12) with the exception that embryo collection was repeated 2-3 times to collect sufficient 1-2 h embryos.

In preparation for microscopic analysis, the methanol was removed, and embryos were treated as described (12). Propidium iodide (PI, Sigma-Aldrich) mounting medium was used for DNA staining, and stained embryos were mounted on glass slides and sealed under a coverslip with nail polish. Imaging was performed with a Zeiss Axioskop microscope with AxioCam MRm camera using 10x and 40x objective lenses.

To assess the cytology of very early-stage embryos from incompatible crosses with *cinB*^*w*No^ or *cinA*-T2A-*cinB*^*w*No^ transgenic males, a slightly different method was employed. Roughly 100 virgin female wCS WT flies and ~100 transgenic *cinB*^*w*No^ or *cinA*-T2A-*cinB*^*w*No^ males were crossed as described previously (12). Embryos were collected from fresh apple juice plates after 30 min, directly dechorionated in 10 mL 50% fresh bleach for 1-3 min, washed once in 10 mL fresh embryo wash buffer (0.6% NaCl, 0.04% Trition X-100) for 5 s, and fixed immediately with methanol or formaldehyde solution. The methanol fixation method (as in the images used for **Figure 5**) (12) and the formaldehyde fixation method (31) were described previously. Embryos were washed and stored in methanol. Embryo collection was repeated 3-4 times to collect sufficient 30-min embryos. All ensuing steps were as above, except the embryos were stained with 40 μL fresh Hoechst 33342 (ThermoFisher Scientific) at 1:1,000 in PBTA buffer. Imaging was done as above using 10x and 100x objective lenses.

### Transgenic fly mRNA expression analysis by RT-qPCR

To analyze expression levels of *w*No *cinA, cinB*, and *cinA-T2A-cinB* transgenes used for hatch-rate assays (**Suppl. Fig. S3**), 11 female or male flies were pooled and kept frozen at −80 °C until processed. Untransformed flies were used as a negative control. Real-time PCR was performed as described previously (23).

### Statistical analysis

All statistical analyses were done with GraphPad Prism (v.9) software. For comparisons between more than two data sets, a non-parametric Kruskal–Wallis one-way ANOVA analysis of variance test followed by Dunn’s multiple comparisons was used in hatch rate analyses. Pairwise *χ*^2^ (Fisher’s exact) test was used for the cytological analyses to infer statistically significant differences between normal and defective cytological phenotypes. A parametric t test was used to compare transgene mRNA levels of transgenic flies.

## Acknowledgements

This study was supported by NIH grant GM136325 to M.H.

## Supplementary Tables

**Table S1.** List of plasmids.

**Table S2.** List of primers. F, Forward, R, Reverse.

## Supplementary Figure legends

**Figure S1**. Growth analysis of yeast W303 cells expressing *cinA*^*w*No^ and/or *cinB*^*w*No^ alleles.

A. Growth assays of yeast expressing *cinA* and *cinB* alleles. W303 cells were transformed with plasmids expressing the indicated alleles from a galactose inducible promoter, and cultures were spotted in serial dilution onto selection plates with the indicated carbon source and grown for 2.5 days at either 30°C, 36°C, or 37°C. EV, empty vector; A, CinA^*w*No^; B, CinB^*w*No^.

B. Expression levels of different CinA and CinB proteins in the same W303 transformants from panel A were measured by immunoblot analysis. Both *Wolbachia* proteins were tagged with a FLAG epitope. PGK (phosphoglycerate kinase), loading control.

**Figure S2**. *w*No CinA and CinB form a cognate-specific protein complex.

GST-pulldown analyses were done with recombinant proteins expressed in separate *E. coli* transformants. Lysates from bacteria expressing the different GST-CinB proteins were first bound to a glutathione resin, The column was washed and then incubated with lysates from cells expressing the indicated CinA proteins. After washing, proteins were eluted with reduced glutathione, resolved by SDS-PAGE, and visualized with GelCode Blue. Cognate protein pairs are shown in lanes 1 and 4. The cartoon interpretation is as in Fig. 3.

**Figure S3**. Relative mRNA levels for *w*No *cinA, cinB*, and *cinA-T2A-cinB* transgenes.

Extracts of transgenic male or female *D. melanogaster* adults were used for RT-qPCR analysis of the indicated transgenes. A compares relative expression of *cinA* in males vs females transgenic for *cinA* alone. B compares relative expression levels of *cinA* in males transgenic for *cinA* alone vs *cinA*-T2A-*cinB*. C compares relative expression levels of *cinB* in males transgenic for *cinB* alone vs *cinA-T2A-cinB*. All quantifications used the 2^−ΔΔCt^ method using *rpl32* as internal reference. **, P<0.01; ****, P< 0.0001 by t test.

