## supplement figures and tables for "The *Wolbachia* CinB Nuclease is Sufficient for Induction of Cytoplasmic Incompatibility"

**Table S1**

| Plasmid name | Description | Tag | |
|---|---|---|---|
| JFB_GA3 | CinA_wNo in PRS425 | N-FLAG | |
| JFB_GB1 | CinB_wNo in pRS416 | N-FLAG | |
| JFB_EB27 | CinA_wPip in PRS425 | N-FLAG | |
| JFB_EA42 | CinB_wPip in pRS416 | N-FLAG | |
| Y16.9.5 | pRS425GAL1 (empty vector) | | |
| Y16.9.6 | pRS416GAL1 (empty vector) | | |
| HC_CN1 | SpeI-CinA_wNo-BamHI in pRS425 | N-FLAG | |
| HC_CO1 | SpeI-CinB_wNo-BamHI in pRS416 | N-FLAG | |
| HC_CinA-wPip | NotI-CinA_wPip-BamHI in pUASp-attb | | |
| GS_005 | BamHI-CinA_wNo-SalI in pET28a-pp | N-His6 | Subclone from HC_CN1 |
| GS_006 | BamHI-CinB_wNo-NotI in pGEX6P1 | N-GST | Subclone from HC_CO1 |
| GS_m004 | BamHI-CinB*(K275A)_wNo-NotI in pGEX6P1 | N-GST | Quickchange from GS_006 |
| GS_m001 | SpeI-CinB*(K275A)_wNo-BamHI in pRS416 | N-FLAG | Quickchange from HC_CO1 |
| GS_m002 | SpeI-CinB*(K629A)_wNo-BamHI in pRS416 | N-FLAG | Quickchange from HC_CO1 |
| MZ | NdeI-CinA_wPip-XhoI In pET28a-pp | N-His6 | Subclone from HC_CinA-wPip |
| JAR_2.2.9 | BamHI-CinB_wPip-XhoI in pGEX6P1 | N-GST | |

**Table S2**

| Primer name | Direction | Sequence (5' - 3') | Restriction site | Description |
|---|---|---|---|---|
| HC369 | F | CGCCTGCATCGAGTGCGTGGAGTCGG | | CinA_wNo Sequencing |
| HC370 | F | CGCCGCGCGGATACGCCCACCTGC | | |
| HC347 | R | GTCCACGGCCTTGTTGAAGCCATCCTCGGCC | | |
| HC319 | R | GCTGCTTCTCCTCGGCGGAGAAGCGGG | | CinB_wNo Sequencing |
| HC320 | F | CCACGACGAGCACGAGGATAGCCTGG | | |
| HC321 | F | GCGAGTTCACCCAGTCCCCACACTGC | | |
| HC343 | F | GCTTCCTGAACAATTGCCCCTCCTTCCTGCAC | | |
| GS003 | F | CGGGATCCATGCCGAAGTCCAAGA | BamHI | CinA_wNo subclone PCR |
| GS004 | R | GCGTCGACTTACTTATTCACCACGGC | SalI | |
| GS008 | F | CGGGATCCATGCACGGCAACAATGAG | BamHI | CinB_wNo subclone PCR |
| GS009 | R | ATTTGCGGCCGCTTAGCGGGAGAAG | NotI | |
| GS014 | F | GCTGGCAGTGGGCGATGAGGAGG | | CinB*(K275A)_wNo Quickchange |
| GS015 | R | CCACTGCCAGCTCGATGATGATCGGC | | |
| GS016 | F | GCTGGCGTACGCCAACACCGCC | | CinB*(K629A)_wNo Quickchange |
| GS017 | R | CGTACGCCAGCTCGATGCCGATGG | | |
| GS022 | F | CCAAGTCGGGAGTGCTGAAT | | CinA_wNo/CinA-T2A-CinB_wNo transgenic fly RT-PCR |
| GS023 | R | GCCAGATTGCGATCCTCCTC | | |
| GS024 | F | GATATCCACGACGAGCACGA | | CinB_wNo/CinA-T2A-CinB_wNo transgenic fly RT-PCR |
| GS025 | R | TCATTGGTCTGATCCACGCC | | |
| GS028 | F | ATGCCAATAGAAACAAAACGTCAGGCTG | | MTD (wMel Wolbachia infection) |
| GS029 | R | CTAGATTCTTCTTTTCTTATGAGAACCAG | | |
| HC373 | F | CAAGTCACTAATCGGTCTTCGAAAGTTCAATATC | | Primers to PCR D.Mel act88F gene |
| HC374 | R | GCACAGCCACGACTCTTACGATTAGTTCTTC | | |
| MZ278 | F | ATGCTAAGCTGTCGCACAAATG | | Drosophila Melanogaster rp49(rpl32) internal reference qPCR |
| MZ279 | R | GTTCGATCCGTAACCGATGT | | |

**Figure S1**

A.

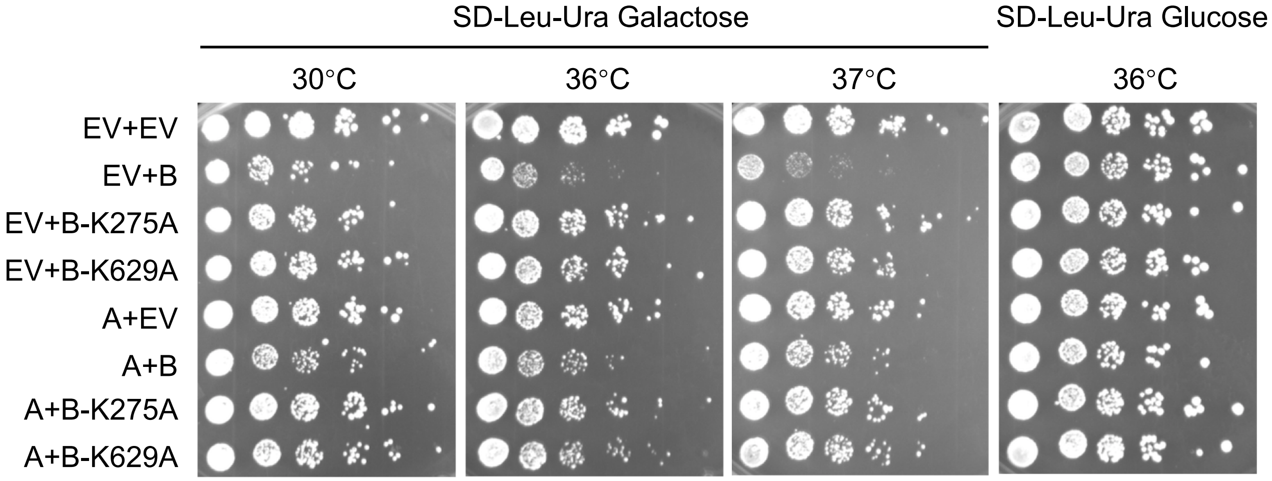


B.

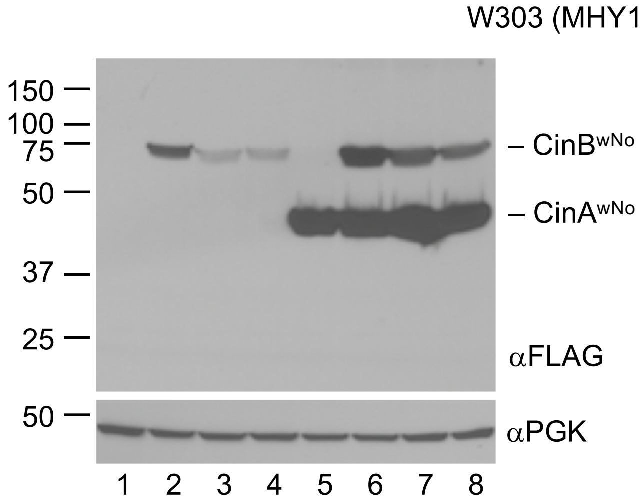


1. pRS416 (EV) + pRS425 (EV)
2. CinB$^{wNo}$ + pRS425
3. CinB$^{wNo}$-K275A + pRS425
4. CinB$^{wNo}$-K629A + pRS425
5. pRS416 + CinA$^{wNo}$
6. CinB$^{wNo}$ + CinA$^{wNo}$
7. CinB$^{wNo}$-K275A + CinA$^{wNo}$
8. CinB$^{wNo}$-K629A + CinA$^{wNo}$

**Figure S2**

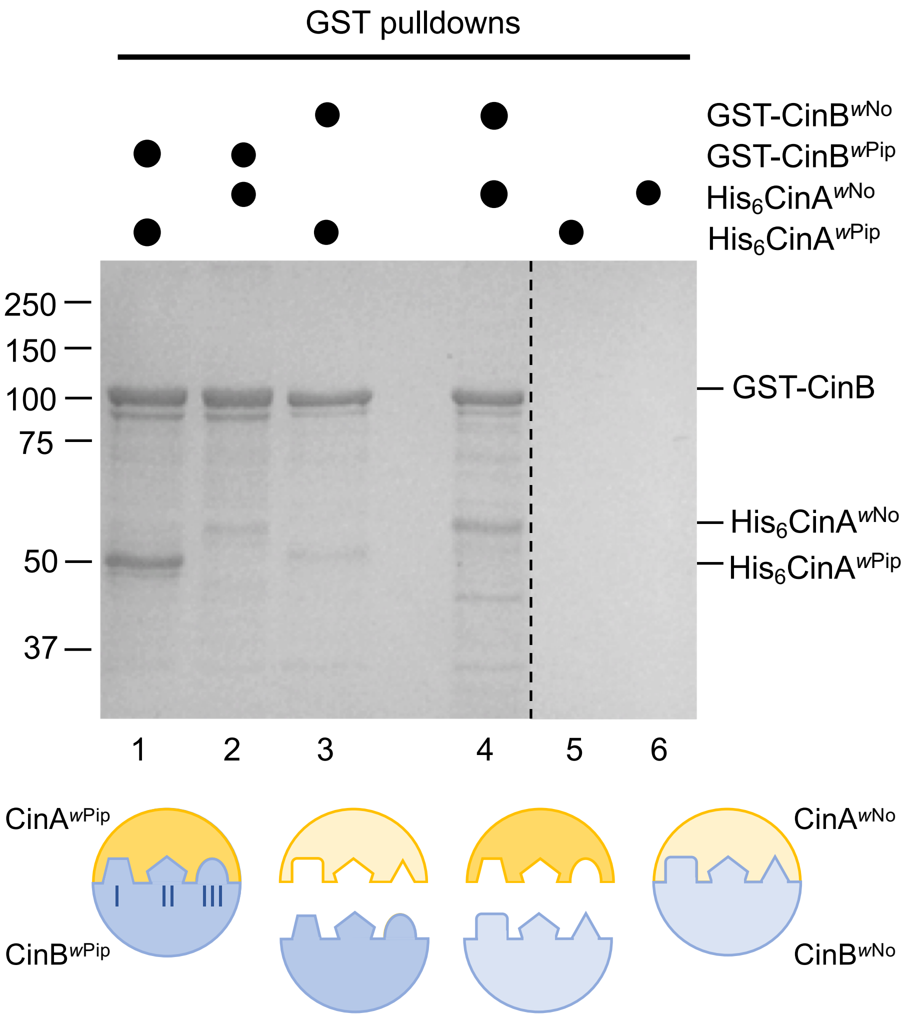

GST pulldowns
GST-CinB$^{wNo}$
GST-CinB$^{wPip}$
His$_6$CinA$^{wNo}$
His$_6$CinA$^{wPip}$
250
150
100
75
50
37
GST-CinB
His$_6$CinA$^{wNo}$
His$_6$CinA$^{wPip}$
1
2
3
4
5
6
CinA$^{wPip}$
CinB$^{wPip}$
I
II
III
CinA$^{wNo}$
CinB$^{wNo}$

**Figure S3**

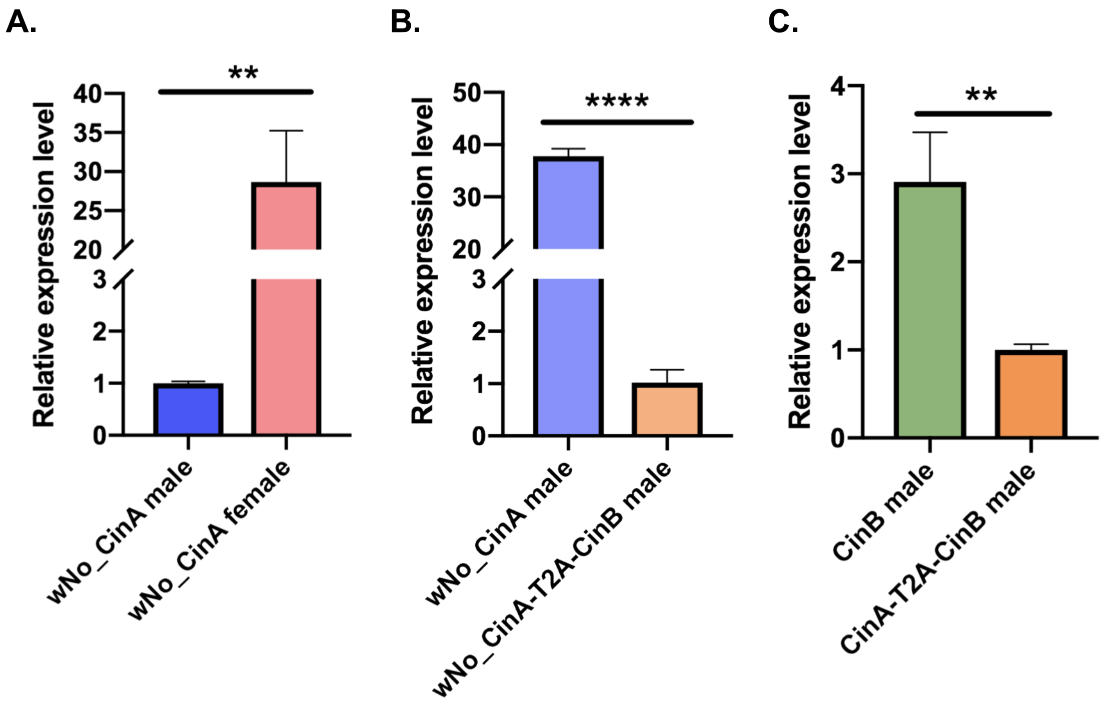

A.
B.
C.
**
****
**
Relative expression level
40
35
30
25
20
3
2
1
0
50
40
30
20
3
2
1
0
4
3
2
1
0
wNo_CinA male
wNo_CinA female
wNo_CinA male
wNo_CinA-T2A-CinB male
CinB male
CinA-T2A-CinB male
